# Investigating the shared genetics of non-syndromic cleft lip/palate and facial morphology

**DOI:** 10.1101/255901

**Authors:** Laurence J Howe, Myoung Keun Lee, Gemma C Sharp, George Davey Smith, Beate St Pourcain, John R Shaffer, Mary L Marazita, Eleanor Feingold, Alexei Zhurov, Evie Stergiakouli, Jonathan Sandy, Stephen Richmond, Seth M Weinberg, Gibran Hemani, Sarah J Lewis

## Abstract

There is increasing evidence that genetic risk variants for non-syndromic cleft lip/palate (nsCL/P) are also associated with normal-range variation in facial morphology. However, previous analyses are mostly limited to candidate SNPs and findings have not been consistently replicated. Here, we used polygenic risk scores (PRS) to test for genetic overlap between nsCL/P and seven biologically relevant facial phenotypes. Where evidence was found of genetic overlap, we used bidirectional Mendelian randomization (MR) to test the hypothesis that genetic liability to nsCL/P is causally related to implicated facial phenotypes. Across 5,804 individuals of European ancestry from two studies, we found strong evidence, using PRS, of genetic overlap between nsCL/P and philtrum width; a 1 S.D. increase in nsCL/P PRS was associated with a 0.10 mm decrease in philtrum width (95% C.I. 0.054, 0.146; P = 0.00002). Follow-up MR analyses supported a causal relationship; genetic variants for nsCL/P homogeneously cause decreased philtrum width. In addition to the primary analysis, we also identified two novel risk loci for philtrum width at 5q22.2 and 7p15.2 in our Genome-wide Association Study (GWAS) of 6,136 individuals. Our results support a liability threshold model of inheritance for nsCL/P, related to abnormalities in development of the philtrum.

## Introduction

Orofacial clefts are malformations characterised by a failure of fusion between adjacent facial structures in the embryo ^1^. Cleft lip with/without cleft palate (CL/P) is a sub-type of orofacial cleft, consisting of individuals presenting with a cleft of the upper lip, with or without a cleft of the palate. Approximately 70% of CL/P cases are non-syndromic, where the facial cleft is not accompanied by other apparent developmental or physical abnormalities ^2^. The non-syndromic form of CL/P(nsCL/P) is a multifactorial trait with both genetic and environmental risk factors ^1^. A possible polygenic threshold model of inheritance is supported by the identification of more than 20 common genetic risk variants for nsCL/P from genome-wide association studies (GWAS) ^3-9^ and single nucleotide polymorphism (SNP) heritability estimates of around 30% ^6^.

Facial morphology in the general population is also likely to be highly polygenic; genome-wide significant loci have been found for multiple facial phenotypes across diverse ethnic populations ^10-14^. In some cases, the genes associated with normal-range variation in facial shape have also been implicated in nsCL/P (e.g. *MAFB*) ^12^. Likewise, previous studies using candidate SNPs have found overlap between nsCL/P risk loci and facial phenotypes in the general population ^11; 15; 16^. For example, the strongest nsCL/P GWAS signal, intergenic variant rs987525 on chromosome 8q24, was found to be associated with more than half of the 48 facial phenotypes studied in a European population ^11^ while in a Han Chinese population, rs642961 in *IRF6* (a major nsCL/P-associated gene) strongly predicted lip-shape variation in females ^16^. However, associations between nsCL/P genetic variants and facial morphology were not consistently replicated, possibly because of methodological differences in measuring facial phenotypes or population differences between cohorts ^10^.

The use of individual markers to demonstrate genetic overlap between two phenotypes has notable limitations; a large number of statistical tests are introduced, and interpretation is difficult when some SNPs show an association and others do not. Polygenic risk scores (PRS) involve incorporating multiple markers, including those not reaching genome-wide significance, into a genetic score that serves as a proxy for a trait ^17^. PRS have been previously shown to be suitable predictors for nsCL/P ^6^ suggesting they can be used to estimate genetic overlap between nsCL/P and normal-range facial morphology.

Interpreting genetic overlap between nsCL/P and a facial phenotype is difficult because the development of the face and development of an orofacial cleft are largely synchronous. One possibility is that differences in the facial phenotype are a sub-phenotypic manifestation of genetic liability to nsCL/P (see Figure 1). The inheritance of dichotomous traits can be modelled on the liability scale, where every individual has an underlying normally distributed liability to the trait determined by genes, environment and chance. Individuals above a liability threshold develop the trait, while increased liability may cause related phenotypic differences in individuals without the trait ^18-20^. For example, increased liability to developing a cleft palate (CP) has been hypothesised to be associated with delayed movement of the palatal shelf, which may in turn result in a CP, dependant on other factors such as shelf and head width ^20^.

**Figure 1.**
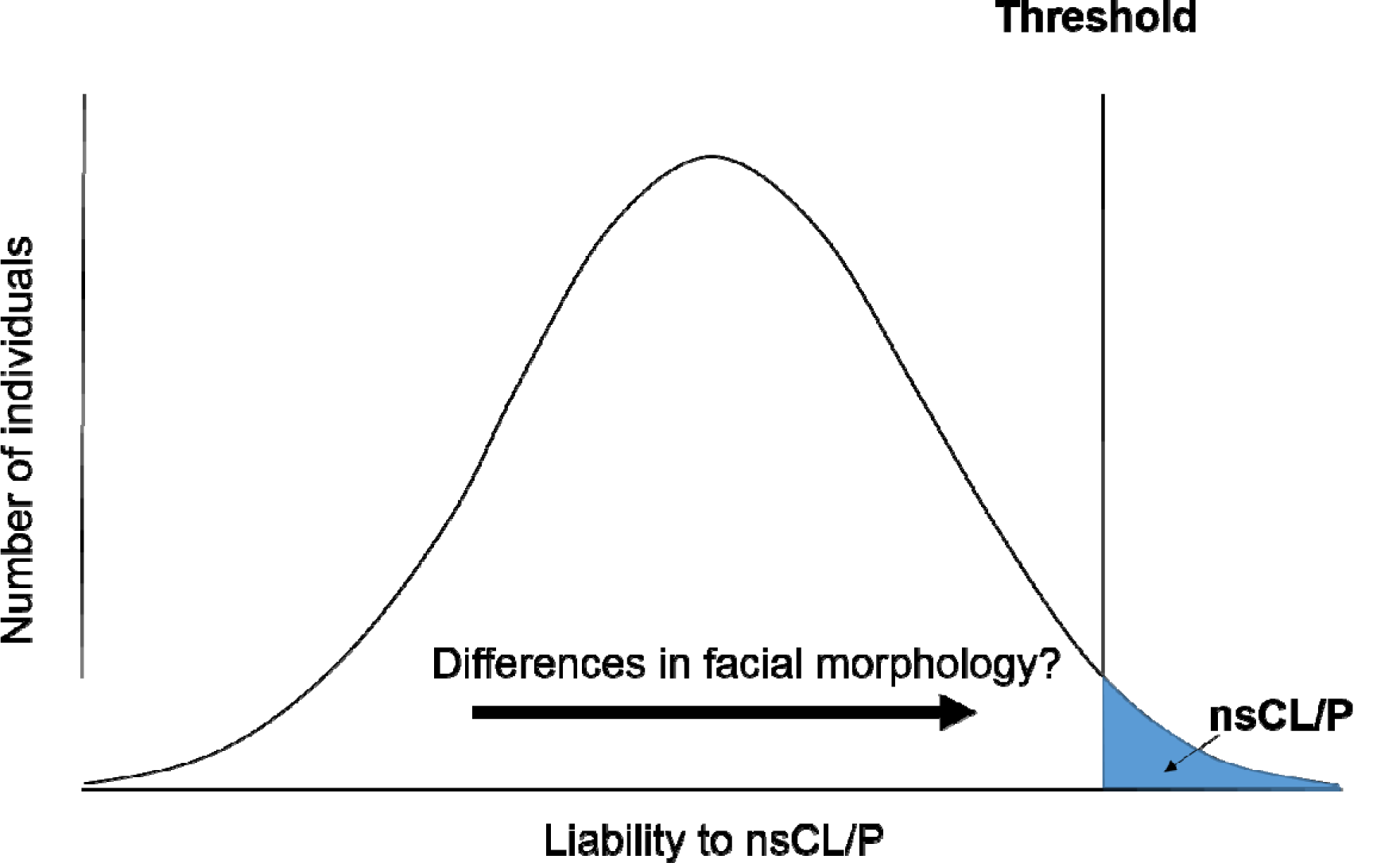
Liability threshold model for nsCL/P. Shown is an illustration of a liability threshold model for nsCL/P. Every individual has a normally distributed liability to nsCL/P, determined by genes, environment and chance. Individuals over the liability threshold develop nsCL/P, with the area under the curve past the threshold equal to the trait incidence. We are hypothesising that liability to nsCL/P, specifically genetic liability to nsCL/P, may be associated with differences in facial morphology across the general population.

In order to evaluate the coherence of the liability-related sub-phenotype model, we apply the principles of Mendelian randomization (MR). MR is an instrumental variable approach, testing causality of an “exposure” and an outcome by using genetic instruments to mimic a randomised controlled trial ^21^. MR relies on several strict assumptions; firstly, genetic variants must be robustly associated with the exposure (in this instance, genetic liability to nsCL/P); secondly, the variants cannot influence the outcome through a pathway independent of the exposure; and thirdly, the variants should not be associated with confounders of the relationship between the exposure and the outcome ^22^. If these assumptions are met, bidirectional MR can be used to test the hypothesis that genetic liability to nsCL/P is causally related to facial morphology ^22^.

In the absence of large-scale publicly available GWAS summary data for nsCL/P, we used individual level genotype data from the International Cleft Consortium to Identify Genes and Interactions Controlling Oral Clefts (ICC) and GWAS summary statistics from the Bonn-II study ^8^ to replicate the meta-analysis GWAS summary statistics from the previously published Ludwig et al 2012 GWAS ^3^. Next, we investigated genetic overlap between nsCL/P and normal-range facial morphology in the general population, using PRS derived from the GWAS summary statistics. Finally, in the instance of genetic overlap, we used bidirectional MR to explore the relationship between nsCL/P and implicated facial phenotypes.

## Subjects and Methods

### Study Participants

#### International Cleft Consortium (ICC)

Data was used on parent-child cleft trios from the ICC (dbGaP Study Accession phs000094.v1.p1) ^23; 24^which includes genotype data from a wide array of geographical locations across North America, Europe and Asia. The data-set included 2,029 parent-offspring trios, 401 parent-offspring pairs, 88 single cleft cases and 25 assorted extended families. Of the 2,543 children with an orofacial cleft; 1,988 presented with nsCL/P while 582 presented with an isolated cleft palate (CPO) and 21 presented with an “unknown cleft”.

Analysis was restricted to trios with a proband diagnosed with nsCL/P. 654 of the parent-offspring trios and 164 of the parent-offspring pairs were of European descent and were used in the primary analyses. 759 parent-offspring trios and 159 parent-offspring pairs of Asian descent were included in secondary analyses.

GWAS genotypes and phenotypes are available at dbGaP (<u>https://www.ncbi.nih/gov/gap;</u> accession number phs000094.v1.p1).

#### ALSPAC

We used data on children from the Avon Longitudinal Study of Parents and Children (ALSPAC), a longitudinal study that recruited pregnant women living in the former county of Avon (UK) with expected delivery dates between 1 April 1991 and 31 December 1992. The initial number of enrolled pregnancies was 14,541, which resulted in 14,062 live births and 13,988 children alive at the age of 1. When the oldest children were approximately 7 years of age, the initial sample was boosted with eligible cases who had failed to join the study originally. For analyses of children after the age of 7, the total possible sample size is 15,247 pregnancies, resulting in 14,775 live births. Full details of the enrolment have been documented elsewhere ^25; 26^. Data have been gathered from the mother and her partner (during pregnancy and post birth) and the children (post birth) from self-report questionnaires and clinical sessions. Ethics approval for the study was obtained from the ALSPAC Ethics and Law Committee and the Local Research Ethics Committee. The study website contains details of all available data through a searchable data dictionary (<u>http://www.bristol.ac.uk/alspac/researchers/dataaccess/datadictionary/</u>).

#### 3D Facial Norms Database

The 3D Facial Norms Database (3DFN) has been described in detail previously ^27^. In brief, we used data from the 3DFN, a database of controls for craniofacial research. 2,454 unrelated individuals of recent European descent, aged between 3 and 40 years were recruited from 4 sites across the USA and screened for a history of craniofacial conditions. 3D-derived anthropometric measurements, 3D facial surface images and genotype data were derived from each study participant.

GWAS genotypes and phenotypes are available at dbGaP (<u>https://www.ncbi.nih/gov/gap;</u> accession number phs000949.v1.p1).

### Facial phenotyping

#### ALSPAC

ALSPAC children were invited to a clinic at the age of 15 years and 5,253 attended, where two high-resolution facial images were taken by Konica Minolta Vivid 900 laser scanners. 4,747 individuals had usable images (506 individuals did not complete the assessment, or the scans were of poor quality and consequently excluded). The coordinates of 22 facial landmarks were derived from the scans. Further methodological details are contained in a previous publication ^10^.

Distances between facial landmarks were computed by calculating the Euclidean distance between the 3D coordinates. To alleviate multiple testing issues, this study chose to test 7 distances that were either tested previously or have biological relevance to nsCL/P (**Supplementary Table 1**). Facial distances used in the analysis are shown in **Figure 2**.

**Figure 2.**
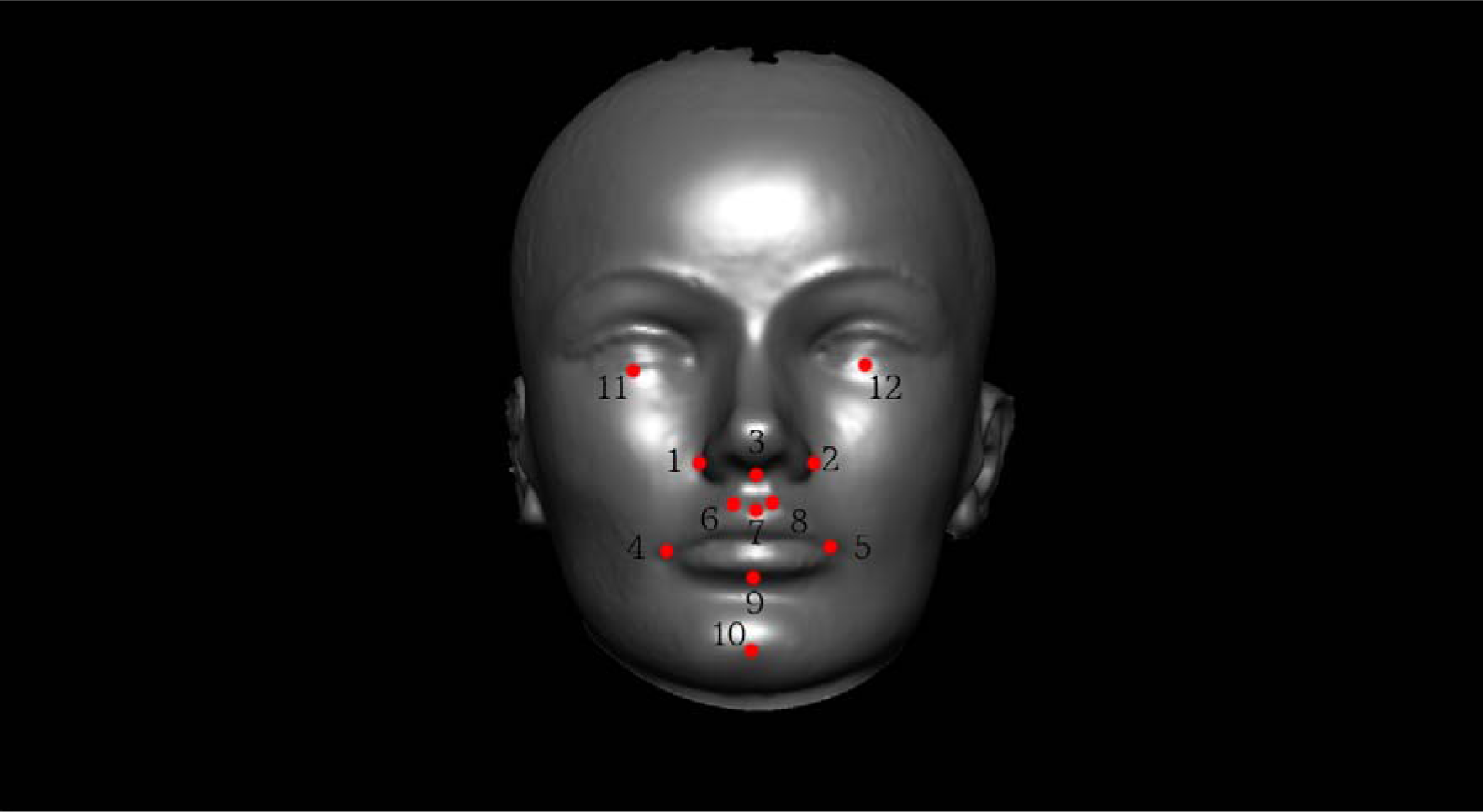
Facial morphological distances of interest. This figure shows the 12 facial landmarks that were used to generate the facial phenotypes tested for association with the nsCL/P PRS. Facial phenotypes were defined as the 3D Euclidean distance between the following landmarks (Nasal width: 1-2, Nasal-lip distance: 3-7, Lip width: 4-5, Philtrum width: 6-8, Lip height: 7-9, Lip-chin distance: 9-10 and inter-palpebrale width: 11-12).

#### 3D Facial Norms Database

A methodological description of the facial phenotyping has been previously described in detail ^27^. In brief, 3DFN study participants had their facial surfaces captured via 3D stereo-photogrammetry using either a two-pod 3dMDface or a multi-pod 3dMDcranial system. Captures were inspected to ensure 3D surface quality and additional captures were obtained if necessary. Similar to ALSPAC, a set of standard facial landmarks were collected from each 3D facial image and linear distances were calculated between the landmark coordinates.

### Genotyping

#### ICC dbGaP

Of 7,347 DNA samples from study subjects genotyped using the Illumina Human610_Quadv1_B array SNP genotyping platform, scans from 7,089 subjects passed QC for unexpected relatedness, gender errors and missingness (>5%) and were released on dbGAP (phs000094.v1.p1). Pre-dbGaP release, SNPs in sample-chromosome combinations with a chromosomal anomaly (e.g. aneuploidy) were excluded. Post dbGaP release, SNPs were excluded for missingness (>5%), MAF (<5%) and HWE (P < 0.05) leaving 490,493 SNPs using PLINK ^28^.

#### ALSPAC

A total of 9,912 ALSPAC children were genotyped using the Illumina HumanHap550 quad genome-wide SNP genotyping platform. Individuals were excluded from further analysis based on having incorrect gender assignments; minimal or excessive heterozygosity (0.345 for the Sanger data and 0.330 for the LabCorp data); disproportionate levels of individual missingness (>3%); evidence of cryptic relatedness (>10% IBD) and being of non-European ancestry (as detected by a multidimensional scaling analysis seeded with HapMap 2 individuals. After quality control for SNPs on missingness (>1%) INFO quality (< 0.8), MAF (< 1%) and HWE (P < 10^-6^) 853,816 SNPs and 8,365 individuals were available for analysis.

#### 3D Facial Norms Database

In collaboration with the Center for Inherited Disease Research (CIDR), 2,454 subjects in the 3DFN database were genotyped using a genome-wide association array including 964,193 SNPs from the Illumina OmniExpress+exome v1.2 array and an additional 4,322 SNPs from previous craniofacial genetic studies. Imputed was performed using the 1000 Genomes reference panel (phase 3) ^27^.

### Statistical analysis nsCL/P meta-analysis Genome-Wide Association Study

The transmission disequilibrium test (TDT) ^29^ evaluates the frequency with which parental alleles are transmitted to affected offspring to test genetic linkage in the presence of genetic association. The TDT was run on 638 parent-offspring trios and 178 parent-offspring duos of European descent to determine genome-wide genetic variation associated with nsCL/P using PLINK ^28^.

The Bonn-II study ^8^ summary statistics from a case-control GWAS of 399 nsCL/P cases and 1,318 controls were meta-analysed, in terms of effect size and standard error, with the TDT GWAS summary statistics using METAL ^30^, based on a previously described protocol for combining TDT and case-control studies ^31^. The final sample consisted of 1215 cases and 2772 parental and unrelated controls.

### Polygenic risk score analysis

#### P-value inclusion threshold determination and PRS construction

We started by estimating the most appropriate P-value inclusion threshold for the nsCL/P PRS. The Bonn-II study summary statistics were used to construct PRS, using different P-value inclusion thresholds, in the independent nsCL/P ICC European and Asian trios (the analysis was done separately for the two ethnic groups). The Polygenic-Transmission Disequilibrium Test (PTDT) ^32^ was then used to measure over-transmission of polygenic risk scores from unaffected parents to affected offspring and thereby select the most predictive P-value inclusion threshold. The P-value inclusion threshold was selected based on the most predictive threshold in the European trios, with results from the Asian trios treated as a sensitivity analysis. Parents with any form of orofacial cleft were removed from this analysis.

Next, using ALSPAC as a reference panel for linkage disequilibrium, PLINK was used to prune and clump the nsCL/P meta-analysis summary statistics (r^2^<0.1 and 250 kb) using the most predictive P-value threshold. The PRS were then constructed in the ALSPAC sample. In addition, power calculations for PRS analysis were performed using AVENGEME ^17; 33^. More information on power calculations is contained in the **Supplementary Methods**.

#### Association of nsCL/P PRS with facial phenotypes in ALSPAC

Of the 4,747 ALSPAC children with face-shape scans, 3,941 individuals had genotype data. GCTA ^34^ was used to prune these individuals for relatedness (IBS < 0.05) and the final sample with complete covariates included 3,707 individuals. The association between facial phenotypes and the nsCL/P PRS was measured in the final sample using a linear regression adjusted for sex, age at clinic visit, height at clinic visit and the first four principal components. Effect sizes were reported per standard deviation increase in PRS.

#### Replication in 3D Facial Norms Database

Distances with some evidence of an association (P < 0.05) in the ALSPAC children were followed up for replication in an independent cohort (3DFN). 2,429 3DFN individuals had genotype and face-shape data. 332 individuals were removed due to missing SNPs in the PRS or missing covariates. The final sample consisted of 2,097 individuals. The association between implicated facial measurements and the nsCL/P PRS was measured using a linear regression adjusted for sex, age, height and the first 4 principal components. Effect sizes were reported per standard deviation increase in PRS.

### Bidirectional Mendelian randomization analysis

A bidirectional two-sample Mendelian randomization analysis was performed using the TwoSampleMR R package ^35^, testing both the forward direction (the effect of genetic risk variants for nsCL/P on implicated facial measurements) and the reverse direction (the effect of genetic risk variants for implicated facial measurements on liability to nsCL/P). The Inverse-Variance Weighted method was used as the primary analysis. Several sensitivity analyses were performed to test the assumptions of MR; the heterogeneity test was used to measure balanced pleiotropy, MR-Egger ^36^ was used to test for directional pleiotropy, the weighted median method ^37^ was used to test if the result is consistent assuming that at least half of the variants are valid and the weighted mode method ^38^ was used to test if the result is consistent assuming that the most common effect is valid. The Steiger test ^39^ was used to determine the likely direction of effect.

#### GWAS summary statistics for nsCL/P and implicated facial phenotypes

MR analysis required relevant SNP association information with respect to both nsCL/P and implicated facial measurements. SNP information relevant to nsCL/P was extracted from the nsCL/P meta-analysis summary statistics, previously described.

For implicated facial phenotypes, GWAS were performed using PLINK ^28^ in both ALSPAC (3,707 individuals) and the 3DFN study (2,429 individuals with genotype and face-shape data), using the same covariates as previously described in the polygenic risk score analysis. Summary statistics were then meta-analysed using METAL ^30^ with the combined sample including 6,136 individuals. SNP information relevant to implicated facial phenotypes was then extracted from the ALSPAC/3DFN meta-analysis summary statistics.

The ALSPAC/3DFN meta-analysis GWAS summary statistics of implicated facial phenotypes were subsequently analysed and functionally annotated ^40^ with the potential overlap between philtrum-width associated SNPs and expression quantitative trait loci (eQTLs) investigated using the Genotype-Tissue Expression (GTEx) catalogue ^41^.

#### Genetic risk variants for nsCL/P and implicated facial phenotypes

For the forward direction, relevant SNPs are variants strongly associated with nsCL/P. 6 well-characterised genome-wide significant nsCL/P SNPs in Europeans were taken from a previous study ^3^.

Information on the nsCL/P SNPs is contained in **Supplementary Table 2**.

For the reverse direction, relevant SNPs are variants strongly associated with the implicated facial phenotypes. We LD clumped (r2<0.001 within 500KB) the ALSPAC/3DFN meta-analysis summary statistics to generate independent instruments for the MR analysis. LD proxies (r2>0.9) were used for SNPs unavailable in the nsCL/P summary statistics and were generated using LDlink and LDproxy ^42^ using the 1000 Genomes CEU/GBR populations as the reference panel ^43^.

### Interpreting bidirectional Mendelian randomization analysis

The results of the bidirectional MR and relevant sensitivity analyses were used to infer the likelihood of the liability-related sub-phenotype model. Three distinct possibilities were considered to explain the association between nsCL/P PRS and implicated facial phenotypes (see Figure 3).

**Figure 3.**
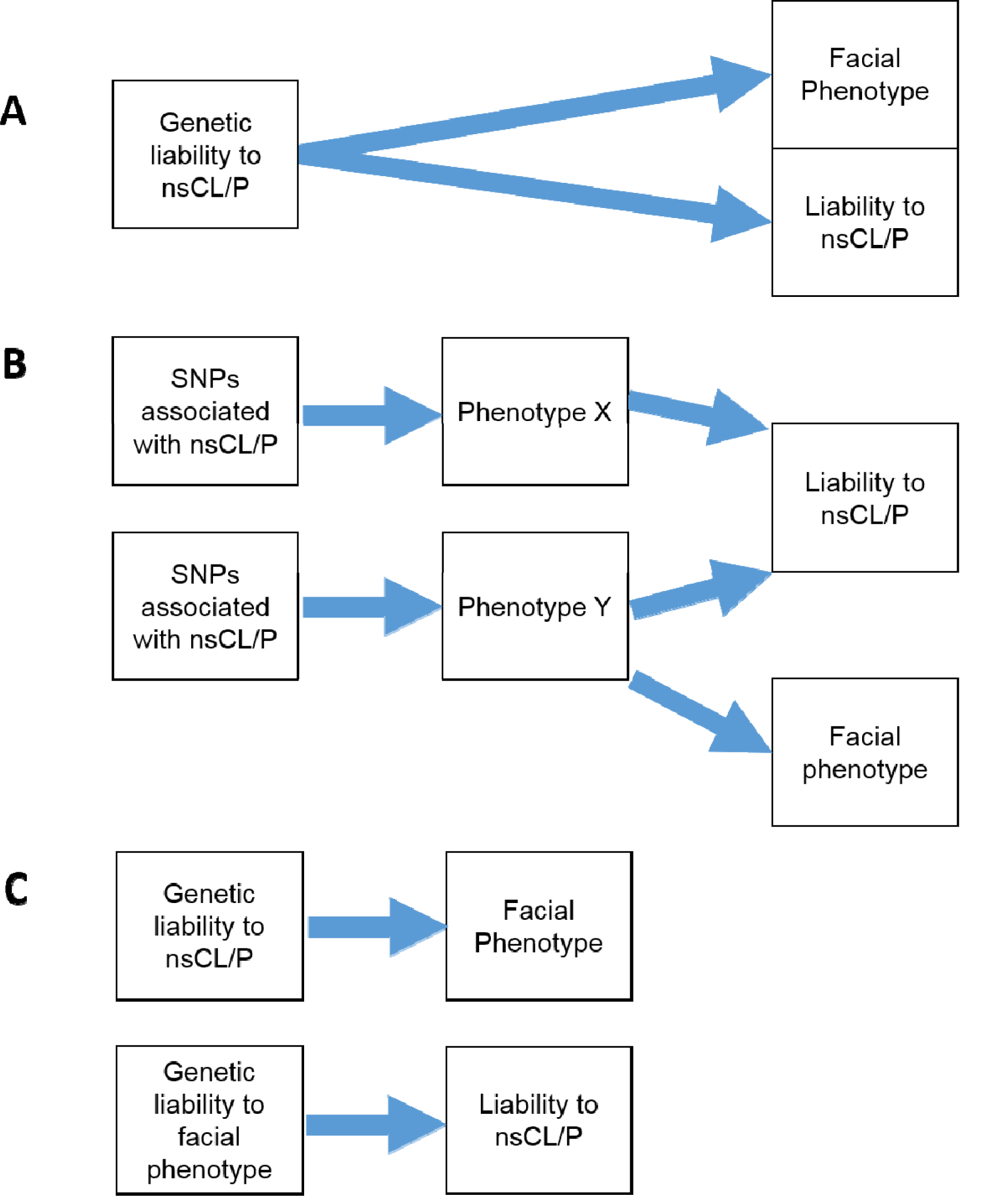
Interpretation of bidirectional MR. (A) *SNPs associated with nsCL/P have a homogeneous effect on the facial phenotype with weak evidence for the reverse direction MR*. We would conclude that genetic liability to nsCL/P causes both increased liability to nsCL/P (in conjunction with the environment and chance) and changes in the facial phenotype. (B) *SNPs associated with nsCL/P have a heterogeneous effect on the facial phenotype*. In this instance, there is weak evidence for genetic liability to nsCL/P causing changes in the facial phenotype because liability assumes a consistent effect. We would conclude that an unknown confounder Y affects the facial phenotype and liability to nsCL/P independently. (C) *SNPs associated with nsCL/P have a homogeneous effect on the facial phenotype AND SNPs associated with the facial phenotype cause increased liability to nsCL/P*. In this instance, there are two possibilities. The first possibility is that the genetic instruments for the facial phenotype are weak (e.g. only one SNP) and so the causal effect estimate of the facial phenotype on liability to nsCL/P is imprecise. The second possibility is that nsCL/P and the facial phenotype have a substantial genetic correlation, which would require further investigation. Here, the results of the Steiger test are useful, as they can infer the most likely direction of effect between nsCL/P and implicated facial phenotypes.

## Results

### Genome-wide association study and genetic proxy for nsCL/P

We performed a GWAS of nsCL/P using the TDT on 638 parent-offspring trios and 178 offspring duos of European descent, and then meta-analysed our results with GWAS summary results previously published on 399 cases and 1317 controls in the Bonn-II study ^8^. This yielded comparable results to a previously published GWAS ^3^, which used a very similar data-set with slightly different quality control and analysis methods (**Supplementary Table 3**).

We also evaluated the predictive accuracy of nsCL/P that could be achieved using different PRS constructed from these summary data by comparing the strength of association at different inclusion thresholds of the PTDT. We determined that including independent SNPs that surpass a P-value threshold of 0.000001 was the most predictive of nsCL/P liability in both European and Asian trios (**Supplementary Table 4**). Therefore, this threshold was used for generating polygenic risk scores from the meta-analysis summary statistics. SNPs included in the selected score are listed in **Supplementary Table 5**.

### The prediction of facial morphology using PRS for nsCL/P

Prior to testing the performance of our nsCL/P PRS on predicting facial morphology, we calculated the minimum genetic correlation required to detect an association between the PRS and the facial phenotypes. We found that the minimum genetic correlation required ranged from 0.17 to 0.28 with differences attributable to different heritability estimates across the facial phenotypes (**Supplementary Table 6**).

We evaluated the performance of our nsCL/P PRS for prediction of seven facial morphological traits. We found evidence of an association between the nsCL/P PRS and philtrum width in the ALSPAC children, where a 1 S.D. increase in nsCL/P PRS was associated with a 0.07 mm decrease in philtrum width (95% C.I. 0.02, 0.13; P=0.014) (**Table 1**).

**Table 1:** Association of nsCL/P PRS with facial phenotypes in ALSPAC children

| 3D facial Euclidean distances in ALSPAC | ALSPAC children (N=3707) |  |
| --- | --- | --- |
|  | Beta (95% C.I.) | P-value |
| Distance between subnasale and labiale superius (Nasal-lip) | -0.25 (-2.16, 1.65) | 0.79 |
| Distance between labiale inferius and pogonion (Lip-chin) | -0.02 (-0.10, 0.06) | 0.64 |
| Distance between left and right palpebrale inferius (Mid-point of eyes) | -0.08 (-0.17, 0.01) | 0.09 |
| Distance between left and right alare (Nasal width) | -0.01 (-0.08, 0.06) | 0.75 |
| Distance between labiales inferius and superius (lip height) | 0.02 (-0.05, 0.10) | 0.53 |
| Distance between left and right crista philtri (philtrum width) | -0.07 (-0.13, -0.02) | 0.014 |
| Distance between left and right cheilion (lip width) | -0.02 (-0.15, 0.10) | 0.70 |

We attempted to replicate this finding in the 3DFN study and found a consistent effect of 1 S.D. increase in nsCL/P PRS being associated with a 0.14 mm decrease in philtrum width (95% C.I. 0.07, 0.21; P = 0.00017). Meta-analysing these results; indicated that a 1 S.D. increase in nsCL/P PRS is associated with a 0.10 mm decrease in philtrum width (95% C.I. 0.054, 0.146; P = 0.00002).

### GWAS of Philtrum width

To generate SNP-philtrum width association information for MR analyses, we performed GWAS of philtrum width in both ALSPAC and 3DFN separately, before meta-analysing. The combined sample included 6,136 individuals of recent European descent. We identified two novel chromosomal regions associated with philtrum width with genome-wide significance at 5q22.2 (lowest P value for rs255877, P=3.8x10^-10^), within the non-coding RNA intronic region of an uncategorised gene *ENSG00000232633*, and 7p15.2 (rs2522825, P=1.4x10^-8^), an intergenic SNP near *HOXA1* (**Supplementary Table 7**). We found some evidence that the two lead SNPs may be eQTLs for nearby genes (**Supplementary Table 8**). The two lead SNPs of the genome-wide significant loci, rs255877 and rs2522825, were used as genetic variants associated with philtrum width in subsequent MR analyses.

### Bidirectional Mendelian randomization

We used MR to investigate the possible causal mechanism that would give rise to the genetic overlap between nsCL/P and philtrum width.

Firstly, we determined whether genetic variants contributing to liability of nsCL/P cause changes in philtrum width, by testing SNPs strongly associated with nsCL/P for association with philtrum width. A 1-unit log odd increase in liability to nsCL/P was associated with a 0.11mm (95% C.I. 0.04, 0.19; P = 0.0036) decrease in philtrum width. Sensitivity analyses suggested there was no evidence of pleiotropy or heterogeneity and validated the consistency of the instrument. Leave-one-SNP-out analysis showed consistent effect estimates after exclusion of each SNP (**Table 2**).

**Table 2:**
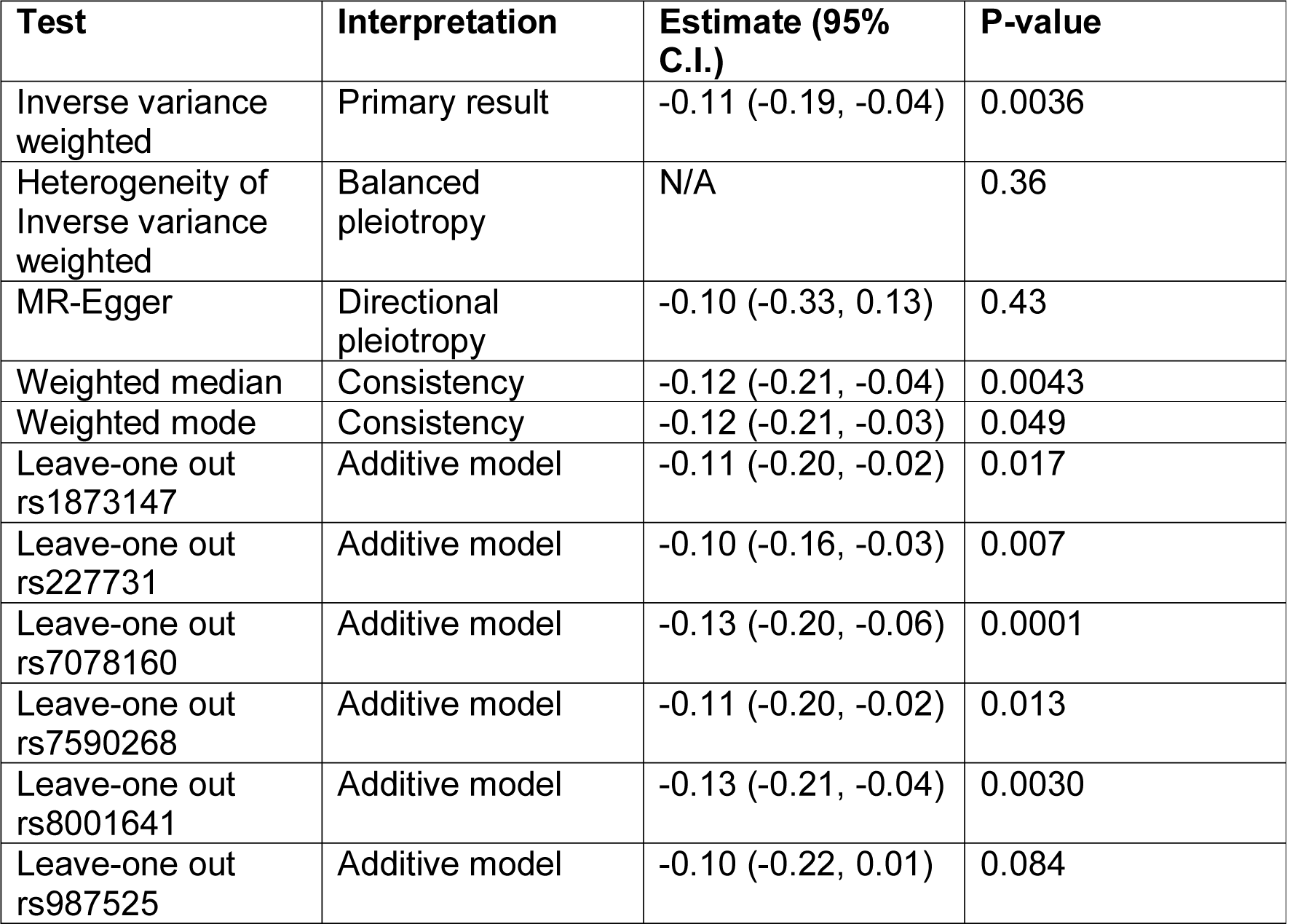
Causal estimates of genetic liability for nsCL/P on philtrum width using Mendelian Randomization

Secondly, we determined whether genetic variants associated with philtrum width also affect liability to nsCL/P, by testing two independent SNPs associated with philtrum width at genome-wide significance (derived in the ALSPAC and 3DFN cohorts) for association with nsCL/P. Utilising strong LD proxies (**Supplementary Table 9**), weak evidence was found of an association between philtrum width-associated variants and liability to nsCL/P (LogOR=0.30; 95% C.I. -0.26, 0.86; P = 0.30). Sensitivity analyses for pleiotropy were limited, with only 2 SNPs.

Thirdly, we used the MR-Steiger test of directionality to test the direction of effect between philtrum width and liability to nsCL/P. The results suggested that the true direction of effect is that genetic variants contributing to liability to nsCL/P cause changes in philtrum width (P <10^-10^).

#### Interpretation of Bidirectional Mendelian randomization

Strong evidence was found for genetic liability to nsCL/P causing decreased philtrum width, weak evidence was found for heterogeneity or assumption violations in the forward-MR, and weak evidence was found for the reverse-MR of philtrum width-associated variants on liability to nsCL/P. Therefore, we conclude that the most likely explanation for the genetic overlap between nsCL/P and philtrum width is that genetic liability to nsCL/P is causally related to decreased philtrum width.

## Discussion

In this manuscript, we have shown that there is genetic overlap between nsCL/P and normal-range variation in philtrum width, and furthermore, that genetic risk SNPs for nsCL/P consistently cause decreased philtrum width in the general population. Notably there was weak evidence for genetic overlap between nsCL/P and upper lip width despite the observational correlation between the widths of the upper lip and philtrum.

There are two main implications of these results. First, our findings demonstrate the aetiological relevance of the formation of the philtrum to nsCL/P. The medial nasal and maxillary processes are responsible for development of the upper lip and philtrum ^44^. Developmental anomalies within these processes may result in a cleft lip ^45^ and our findings show that even when there is successful fusion, as in our study populations, the genetic variants which give rise to a CL/P cause decreased philtrum width. Secondly, the non-heterogenous additive effect of common nsCL/P risk variants, on a related phenotype in the general population, supports a polygenic threshold model of inheritance for nsCL/P.

Although previous studies have looked at nsCL/P related sub-phenotypes, this study uses causal inference methods to more formally investigate the relationship. Our identification of phenotypic differences related to nsCL/P liability are consistent with previous studies ^46-51^ observing sub-clinical facial phenotypes in individuals with nsCL/P and their unaffected family members, particularly a previous study which observed reduced philtrum width in unaffected parents of individuals with nsCL/P ^51^. A polygenic threshold model of inheritance related to development of the philtrum is consistent with a previously proposed mechanism for the inheritance of cleft palate ^20^, the identification of numerous common nsCL/P genetic risk variants ^3-7^ and estimation of a substantial SNP heritability for nsCL/P ^6^. We do not replicate associations between nsCL/P and other facial morphological dimensions found in previous studies ^11; 15; 51^ using candidate SNPs but note that polygenic risk score methods are methodologically distinct and are used to investigate a different research question to single SNP analyses.

We extend the investigation of the association between nsCL/P and facial morphology in two important ways. We demonstrate that the association is present not only in unaffected family members but also in the general population, and use MR to demonstrate that this relationship is present on the liability scale. Conventionally MR is used to test possible causal effects of a modifiable continuous exposure such as cholesterol or alcohol on disease outcomes ^52; 53^. Here we exploit the principles of MR to test the threshold hypothesis, by inferring a causal relationship between genetic variants contributing to liability of nsCL/P and philtrum width in a non-clinical population. We interpret this causal relationship as evidence that smaller philtrum width is a sub-phenotypic manifestation attributable to the same genetic variants that cause nsCL/P.

In addition to investigating the relationship between facial morphology and nsCL/P, we also performed the first GWAS of philtrum width, and identified two novel genome-wide significant loci. We found some evidence of an overlap between philtrum width associated loci and eQTLs that warrants further investigation.

The causal inference made in this study was achieved through the use of two independent cohorts as discovery and replication samples which greatly reduces the risk of false positives and demonstrates that results can be generalised to different populations. Detailed facial phenotyping data on a large number of individuals in our cohorts along with other detailed phenotype and genotype data enabled us to identify philtrum width as being the most relevant facial morphological feature from amongst seven biologically likely candidates. Statistical power does limit the detection of other features that may have mechanistic relationships with smaller effect sizes (**Supplementary Table 6**).

In this study, we combined CL/P and cleft lip only (CLO), however there is evidence suggesting that there are distinct aetiological differences between these traits, ^5; 54; 55^ which could reduce our statistical power, and complicates interpretation. For example, the philtrum may be more related to CLO, but we did not have sufficient data to compare nsCL/P subtype differences. An additional limitation is that there are few well-characterised genetic risk loci for philtrum width, so our MR analysis testing if genetic variants associated with a narrow philtrum width also affect liability of nsCL/P, may be underpowered.

We conclude that genetic liability to nsCL/P is causally related to variation in philtrum width and that this finding supports a polygenic threshold model of inheritance for nsCL/P, related to abnormalities in development of the philtrum. Further research looking at the relationship between genetic liability for nsCL/P and severity of cleft would provide further evidence for the polygenic threshold model.

## Acknowledgements

We are extremely grateful to all the families who took part in the ICC, ALSPAC and 3DFN studies as well as the many individuals involved in the running of the studies and recruitment, which include midwives, interviewers, computer and laboratory technicians, clerical workers, research scientists, volunteers, managers, receptionists, and nurses.

Funding support for the ICC study was provided by several previous grants from the National Institute of Dental and Craniofacial Research (NIDCR). Funding for individual investigators include: R21-DE-013707 and R01-DE-014581 (Beaty); R37-DE-08559 and P50-DE-016215 (Murray, Marazita) and the Iowa Comprehensive Program to Investigate Craniofacial and Dental Anomalies (Murray); R01-DE-09886 (Marazita), R01-DE-012472 (Marazita), R01-DE-014677 (Marazita), R01-DE-016148 (Marazita), R21-DE016930 (Marazita); R01-DE-013939 (Scott). Parts of this research were supported in part by the Intramural Research Program of the NIH, National Institute of Environmental Health Sciences (Wilcox, Lie) and additional recruitment was supported by the Smile Train Foundation for recruitment in China (Jabs, Beaty, Shi) and a grant from the Korean government (Jee). The genome-wide association study, also known the Cleft Consortium, is part of the Gene Environment Association Studies (GENEVA) program of the trans-NIH Genes, Environment and Health Initiative [GEI] supported by U01-DE-018993. Genotyping services were provided by the Center for Inherited Disease Research (CIDR). CIDR is fully funded through a federal contract from the National Institutes of Health (NIH) to The Johns

Hopkins University, contract number HHSN268200782096C. Funds for genotyping were provided by the NIDCR through CIDR’s NIH contract. Assistance with genotype cleaning, as well as with general study coordination, was provided by the GENEVA Coordinating Center (U01-HG-004446) and by the National Center for Biotechnology Information (NCBI). Datasets used for the analyses described in this manuscript were obtained from dbGaP at [https://www.ncbi.nlm.nih.gov/projects/gap/cgi-bin/study.cgi?study_id=phs000094.v1.p1] through dbGaP accession number [phs000094.v1.p1].

The analysis performed in this study was supported by the UK Medical Research Council (MC_UU_12013) and the Cleft Collective funded by the Scar Free Foundation (REC approval 13/SW/0064). The UK Medical Research Council and the Wellcome Trust (Grant ref: 102215/2/13/2) and the University of Bristol provide core support for ALSPAC. The National Institute for Dental and Craniofacial Research (<u>http://www.nidcr.nih.gov/</u>) provided funding, for 3DFN related analyses, through the following grants: U01-DE020078; R01-DE027023; R01-DE016148. The funders had no role in study design, data collection and analysis, decision to publish, or preparation of the manuscript.

